# Combined multiphoton microscopy and somatostatin receptor type 2 imaging of pancreatic neuroendocrine tumors

**DOI:** 10.1101/2023.02.03.526958

**Authors:** Noelle Daigle, Thomas Knapp, Suzann Duan, David W. Jones, Ali Azhdarinia, Sukhen C. Ghosh, Solmaz AghaAmiri, Naruhiko Ikoma, Jeannelyn Estrella, Martin J. Schnermann, Juanita L. Merchant, Travis W. Sawyer

## Abstract

Pancreatic neuroendocrine tumors (PNETs) are a rare but increasingly more prevalent cancer with heterogeneous clinical and pathological presentation. Surgery is the preferred treatment for all hormone-expressing PNETs and any PNET greater than 2 cm, but difficulties arise when tumors are multifocal, metastatic, or small in size due to lack of effective surgical localization. Existing techniques such as intraoperative ultrasound provide poor contrast and resolution, resulting in low sensitivity for such tumors.

Somatostatin receptor type 2 (SSTR2) is commonly overexpressed in PNETs and presents an avenue for targeted tumor localization. SSTR2 is often used for pre-operative imaging and therapeutic treatment, with recent studies demonstrating that somatostatin receptor imaging (SRI) can be applied in radioguided surgery to aid in removal of metastatic lymph nodes and achieving negative surgical margins. However not all PNETs express SSTR2, indicating labeled SRI could benefit from using a supplemental label-free technique such as multiphoton microscopy (MPM), which has proven useful in improving the accuracy of diagnosing more common exocrine pancreatic cancers.

Our work tests the suitability of combined SRI and MPM for localizing PNETs by imaging and comparing samples of PNETs and normal pancreatic tissue. Specimens were labeled with a novel SSTR2-targeted contrast agent and imaged using fluorescence microscopy, and subsequently imaged using MPM to collect four autofluorescent channels and second harmonic generation. Our results show that a combination of both SRI and MPM provides enhanced contrast and sensitivity for localizing diseased tissue, suggesting that this approach could be a valuable clinical tool for surgical localization and treatment of PNETs.

## 1. INTRODUCTION

Pancreatic neuroendocrine tumors (PNETs) are a heterogeneous disease with a low 5-year survival rate of 27-38%.^1^ Improved surveillance and imaging in recent years has increased the number of PNET cases being diagnosed, and has led to a decrease in the average tumor size at diagnosis. As a result, incidence of PNETs has surged six-fold over the past two decades, making it an increasing clinical problem. Surgery is the preferred method of treatment for the majority of PNETs >= 2 cm and any presenting with clinical symptoms, but current surgical localization techniques such as ultrasound provide poor contrast and resolution, ultimately increasing the risk of positive margins and incomplete resections;^2, 3^ the National Comprehensive Cancer Network reported the global disease recurrence for PNETS to be between 21-42% in 2018.^2^ Furthermore, resection is particularly difficult for small or multifocal PNETs, prompting surgeons to perform more demolitive resections than are necessary, increasing patient mortality and reducing quality of life.^2, 3^

Multiphoton microscopy (MPM) is a fast-growing optical imaging technique with greater tissue penetration depth and reduced out-of-plane photodamage compared to single photon microscopy. One subfield, label-free MPM, does not rely on antibody labeling or fluorescent markers, instead using photophysical processes to image naturally occurring tissue biomarkers. This technique can notably collect autofluorescence and second harmonic generation (SHG), making MPM well suited for biomedical imaging as numerous autofluorescent metabolites exist in human tissue, such as nicotinamide adenine dinucleotide and hydrogen (NADH), flavin adenine dinucleotide (FAD), lipofuscins, and porphyrins.^4^ NADH and FAD together inform cellular redox reactions and metabolism, while lipofuscins serve as a marker of cellular oxidation and senescence, and porphyrins denote the level of tissue vascularization.^5^ These autofluorescent molecules can be used to provide a measurement of cellular activity and their atypical expression has been reported in cases of cancer;^6^ SHG is a nonlinear, non-fluorescent light scattering event exhibited by non-centrosymmetric or hyperpolarizable molecules such as collagen, which can be used to identify connective tissues.^4^ However, MPM is limited by its narrow field of view, indicating combining this technology with a wide-field imaging modality would be beneficial for large-scale tissue imaging.

Somatostatin is a growth hormone-inhibiting peptide produced by delta cells in the pancreas. There are five somatostatin receptor subtypes labeled 1-5, also expressed within the pancreas.^7^ Notably, somatostatin receptor type 2 (SSTR2) is overexpressed in over 80% of PNETs.^2^ SSTR2 is often used for pre-operative imaging and therapeutic treatment,^3^ with recent studies demonstrating that somatostatin receptor imaging (SRI) can be applied in radioguided surgery to aid in removal of metastatic lymph nodes and achieving negative surgical margins.^8^ This suggests that fluorescently-labeled SSTR2 imaging could be used in conjunction with MPM for wide-field localization and complement the inherent small field of view of MPM.

Low overall survival, small average tumor size, poor current localization techniques, and the inherent complications of pancreatic surgery all make PNETs a compelling choice for applying MPM during surgical intervention for microscopic tumor localization and margin definition. Here we test the suitability of such a concept by imaging patient-derived PNETs and normal pancreatic tissue using both MPM and SRI, and training a machine learning algorithm to classify the two tissue types using this data. Ultimately, this algorithm will be utilized to provide a spatial mapping of the probability of a tissue being diseased, permitting surgical guidance in real time.

## 2. METHODS

A set of 12 formalin-fixed frozen patient samples comprised of six PNETs and six normal pancreas tissue were obtained from the University of Arizona Tissue Acquisition and Cellular/Molecular Analysis Shared Resource. Samples were embedded in Optimal Cutting Temperature (OCT) compound and cryosectioned in five-micron slices. Tumor and normal samples originated from the same patient in three cases. However, this was not possible for all cases as the relative rarity of this cancer limits availability of samples. All samples were de-identified to protect patient confidentiality.

### 2.1 Somatostatin Receptor Imaging

All 12 samples were stained with a novel fluorescent near-infrared SSTR2-targeted peptide, MMC(FNIR-Tag)-TOC, developed by the Azhdarinia laboratory following their established protocol,^9^ and subsequently wet mounted using Fluoromount-G®. Samples were then imaged with an inverted Olympus IX71 microscope with a 20x objective (1-U2B825-U, Olympus), illuminated with a xenon light source (U-LH75XEAPO, Olympus), and and ORCA-Flash 4.0 digital CMOS camera (C11440-22CU, Hamamatsu Photonics K.K.). Images were collected as 2048 x 2048 pixel tiles at 16-bit depth using a manual stage with varying overlap.

### 2.2 Multiphoton Microscopy

Five wavelength channels were obtained using MPM, selected to probe four endogenous fluorophores that are common biomarkers of disease (FAD, NADH, lipofuscin, and porphyrin) as well as SHG. These were selected based on previous work to image neuroendocrine tumors, where these markers demonstrated high contrast to disease.^6^ Table 1 shows the excitation and emission wavelengths used for each channel. The four fluorophore channels have some overlap, which we reduced by choosing excitation wavelengths to minimize emission wavelength range overlap. However, it is important to note that while these channels contain signal emitted by the four fluorophores, not all signal in each channel is specific to each fluorophore (e.g., autofluorescence produced by other biological molecules may contribute to the signal intensity of each spectra, such as the broad fluorescence emission by collagen).

**Table 1.** Excitation and emission wavelengths used for each of the five MPM channels.

| Dominant Channel Fluorophore | Excitation $\lambda$ [nm] | Emission $\lambda$ [nm] |
| --- | --- | --- |
| NAD(P)H | 750 | 425–465 |
| FAD | 920 | 475–600 |
| Lipofuscins | 830 | 550–600 |
| Porphyrins | 900 | 600–650 |
| SHG | 880 | 430–450 |

Images were collected using the Zeiss LSM880 NLO upright multiphoton microscope with a 20x objective in the Optical Imaging Core at the University of Arizona with a tunable laser and detector. Regions were selected on each sample to cover the same area as the SSTR2 fluorescence images. Large gray-scale images were acquired using an automatic motorized stage as a grid of 13 x 13 tiles with 10% overlap. Tiles were 256 x 256 pixels with 16-bit depth. To account for tilt or unevenly sliced samples, images were taken in five z-stacks at 2 micron intervals. The whole image was assembled from tiles into stitched mosaics using several artifact correction techniques outlined in Figure 1. For a full discussion of image reconstruction methods used in this work, please refer to Knapp et al.^10^ Whole images were typically 3016 x 3016 pixels, covering an area of roughly 4 mm x 4 mm on the sample, with the exception of one 2326 x 3706 pixel sample (due to sample geometry needing a different aspect ratio).

**Figure 1.**
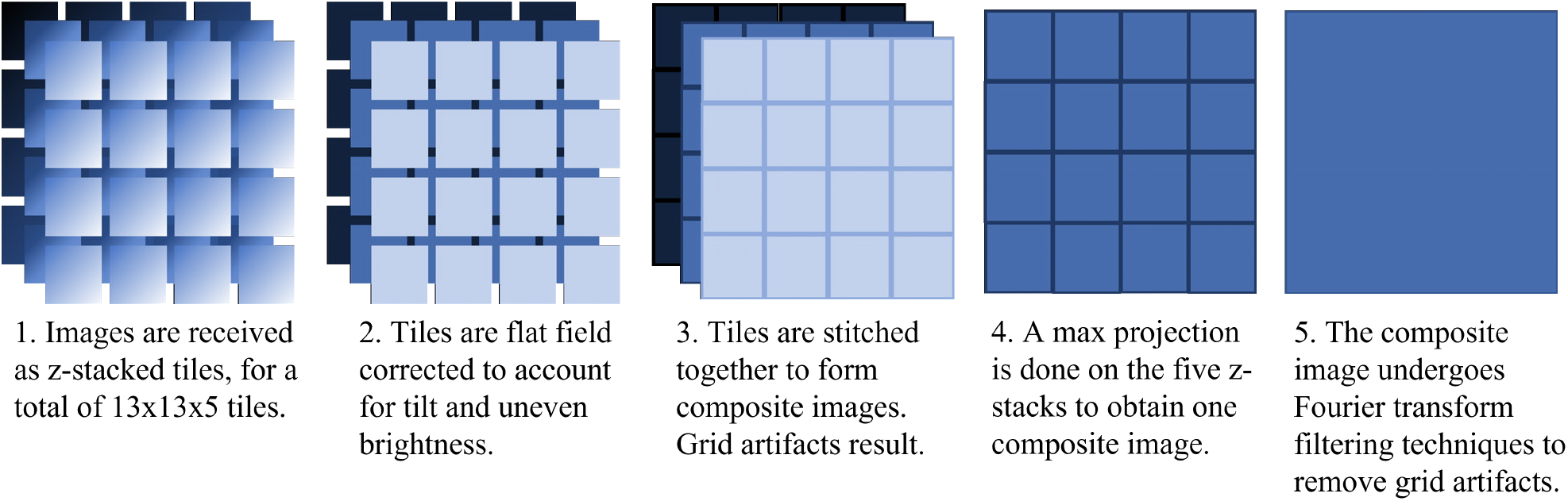
Methods used to stitch and artifact-correct MPM images.^10^ In step 3, tiles were stitching using the ImageJ Grid/Collection stitching plugin.^11^

### 2.3 Image Analysis and Classification

A total of six images were acquired for each patient sample: four autofluorescent channels, one SHG channel, and one SSTR2 fluorescence image. First, a lower threshold applied was applied to remove background and noise, which was determined by calculating the average value in a region of the image with no tissue. The FAD image required an upper threshold applied as well in order to mask out lipofuscin deposits in this channel. Threshold values are available in Table 2.

**Table 2.** 16 bit pixel value threshold levels for all MPM images.

| Channel | Lower Threshold [pixel value] | Upper Threshold [pixel value] |
| --- | --- | --- |
| NAD(P)H | 7000 | 65535 |
| FAD | 3000 | 50000 |
| Lipofuscins | 3000 | 65535 |
| Porphyryns | 3000 | 65535 |
| SHG | 3000 | 65535 |
| SSTR2 | 1500 | 65535 |

13 texture features, listed in Figure 2, were calculated for each of the four autofluorescent and SHG images using Haralick’s method for feature extraction. This method is based off of evaluation of a gray level co-occurrence matrix (GLCM). These matrices describe the relationship of two pixels on the image with each other and are based on the spatial relationship of the two pixels in question, specified by a distance and an angle, and the gray tones of the pixels. Each pixel in the image, excluding the very outer pixels, has eight immediate neighbors. Therefore, there are four possible angles between each pixel and its neighbors: 0, 45, 90, and 135 degrees. For each pixel distance, four GLCM can be calculated for each of the four unique angles.^12^ Our analysis used a pixel distance of 1 and averaged the four GLCM in each direction as the sample can be treated as rotationally symmetric.

**Figure 2.**
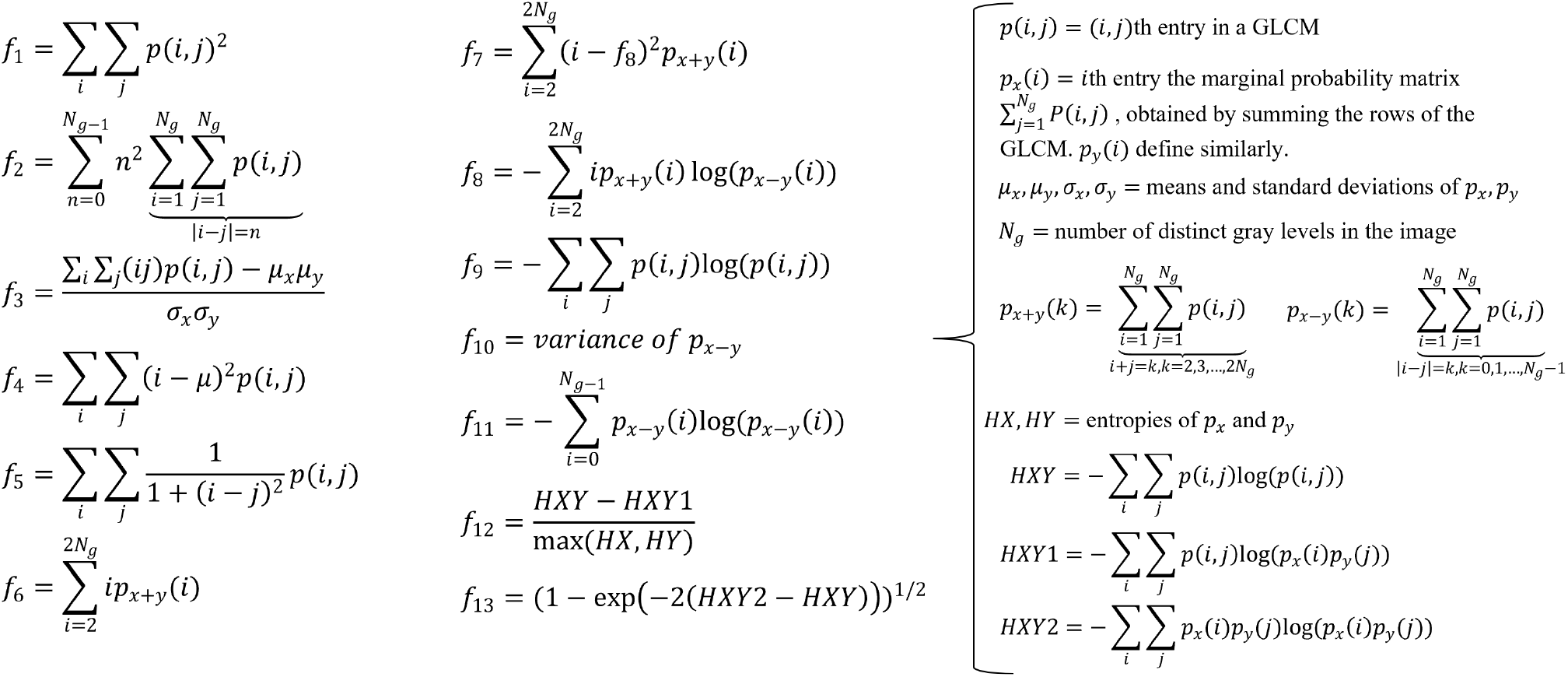
The 13 texture features used to develop classifiers.^12^ Features are labeled *f*_1_ through *f*_13_. Supplemental equations used to calculate the features are listed in the right column of the figure.

For each patient sample, a set of ten tiles were selected as regions of interest from the SSTR2 fluorescence image and the average pixel intensity was calculated to simulate raw fluorescence signal (as would be observed in a wide-field surgical localization paradigm). In total there are 66 features for each sample, 65 from MPM and one from SSTR2 imaging.

Linear discriminant analysis (LDA) is a supervised analysis method that proposes to use a linear combination of features to reduce dimension. It finds projections that maximize the differences between classes while minimizing differences within each class.^13^ In our study we investigated classifiers developed using optimal features determined by LDA using singular value decomposition in order to ascertain the lowest number of features needed to classify tumor and normal tissue to highest accuracy. We tested sets of features ranging from 1 to 4 features. The models were validated using a leave-one-out-approach. Classification accuracy and receiver operator characteristic (ROC) curves were recorded.

## 3. RESULTS AND DISCUSSION

### 3.1 Image Features

Figure 3 shows all five MPM and one example SSTR2 fluorescence image for one tumor and one normal sample. Also shown is a histological reference image used to establish a ground truth tissue status.

**Figure 3.**
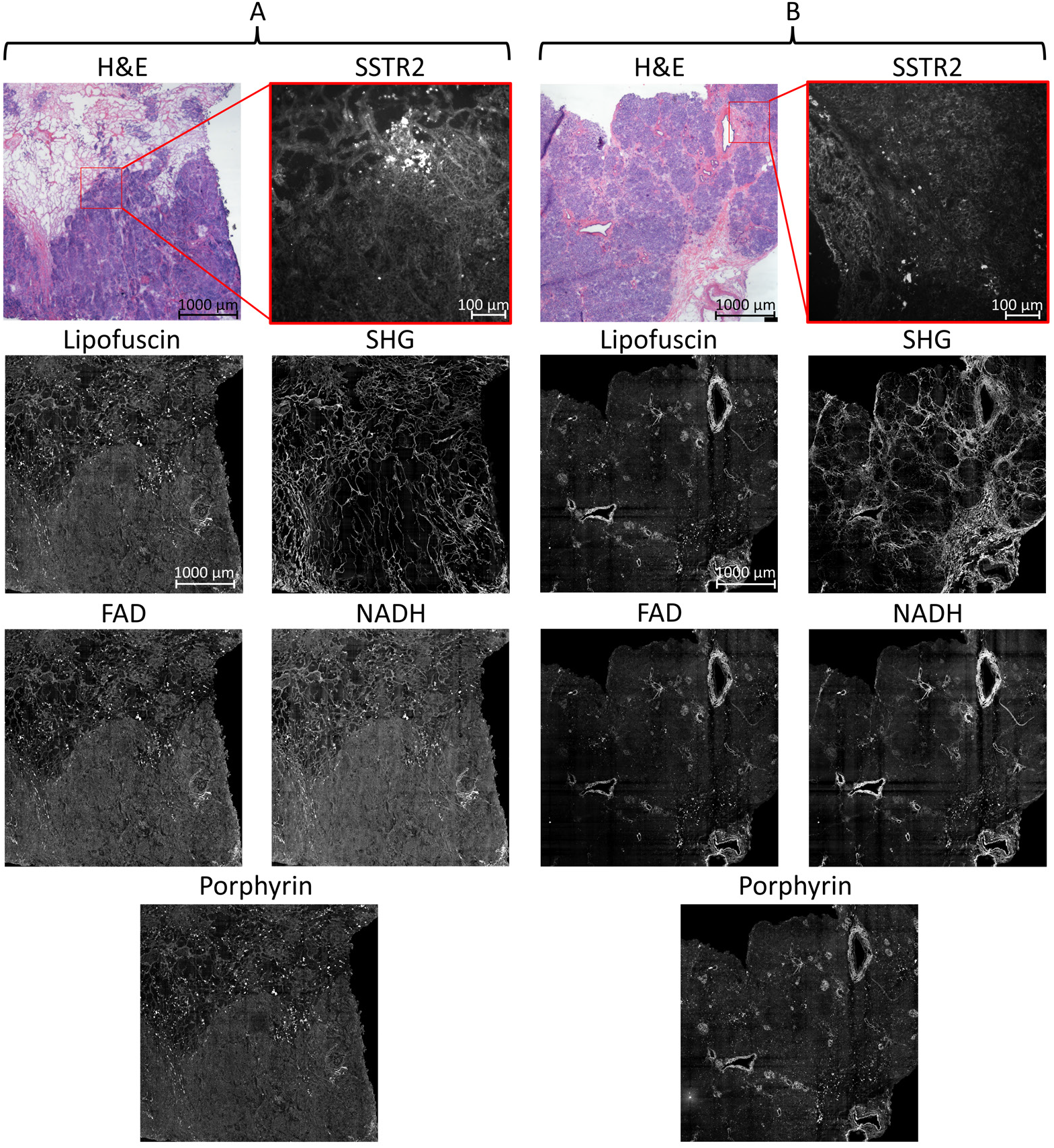
All five MPM images, one selected SSTR2 fluoresence region of interest tile, and a hematoxylin and eosin (H&E) stained slide for (A) a tumor and (B) a normal sample. Images were uniformly artificially brightened for qualitative observation of imaging features.

Figure 4 is a heatmap visualizing correlation between imaging channels averaged across all samples. We note a low correlation between SHG and the four autofluorescent MPM channels as expected, as SHG is a light-scattering event distinct from fluorescence. We see moderate correlation between the four autofluorescent MPM channels, although Lipofuscin has higher correlation with the other channels than FAD, NADH, or porphyrin do, due to its wide range of emission wavelengths. This figure motivates the inclusion of more than one MPM channel in order to capture more information within the dataset.

**Figure 4.**
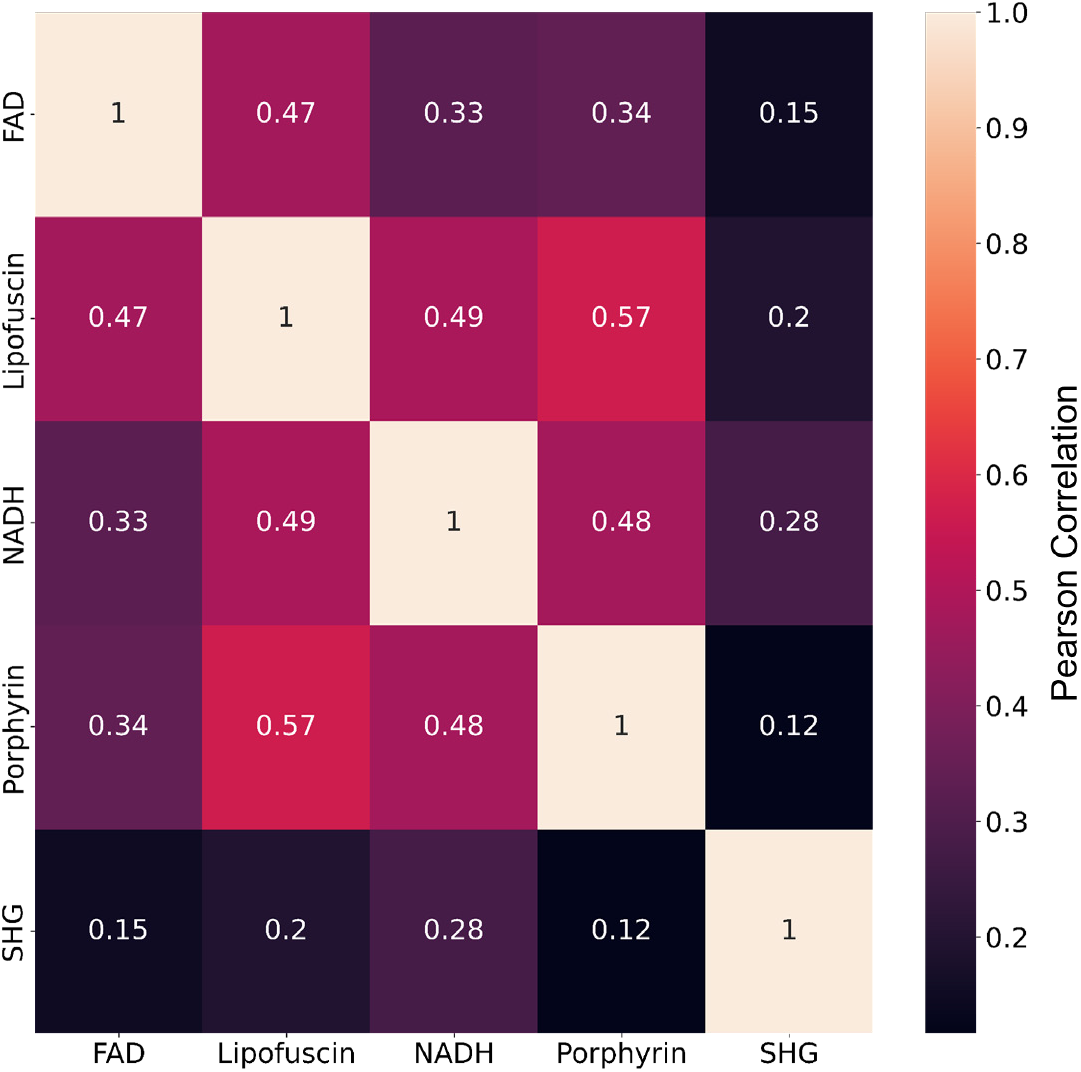
Heat matrices showing correlation between imaging channels averaged across all samples.

Figure 5 shows Z-scores for each of the 66 features for all samples. We see a general trend of normal samples having low Z-scores when tumor samples have high Z-scores, indicating there are distinct qualities that describe each tissue type that Haralick’s feature extraction is able to quantify.

**Figure 5.**
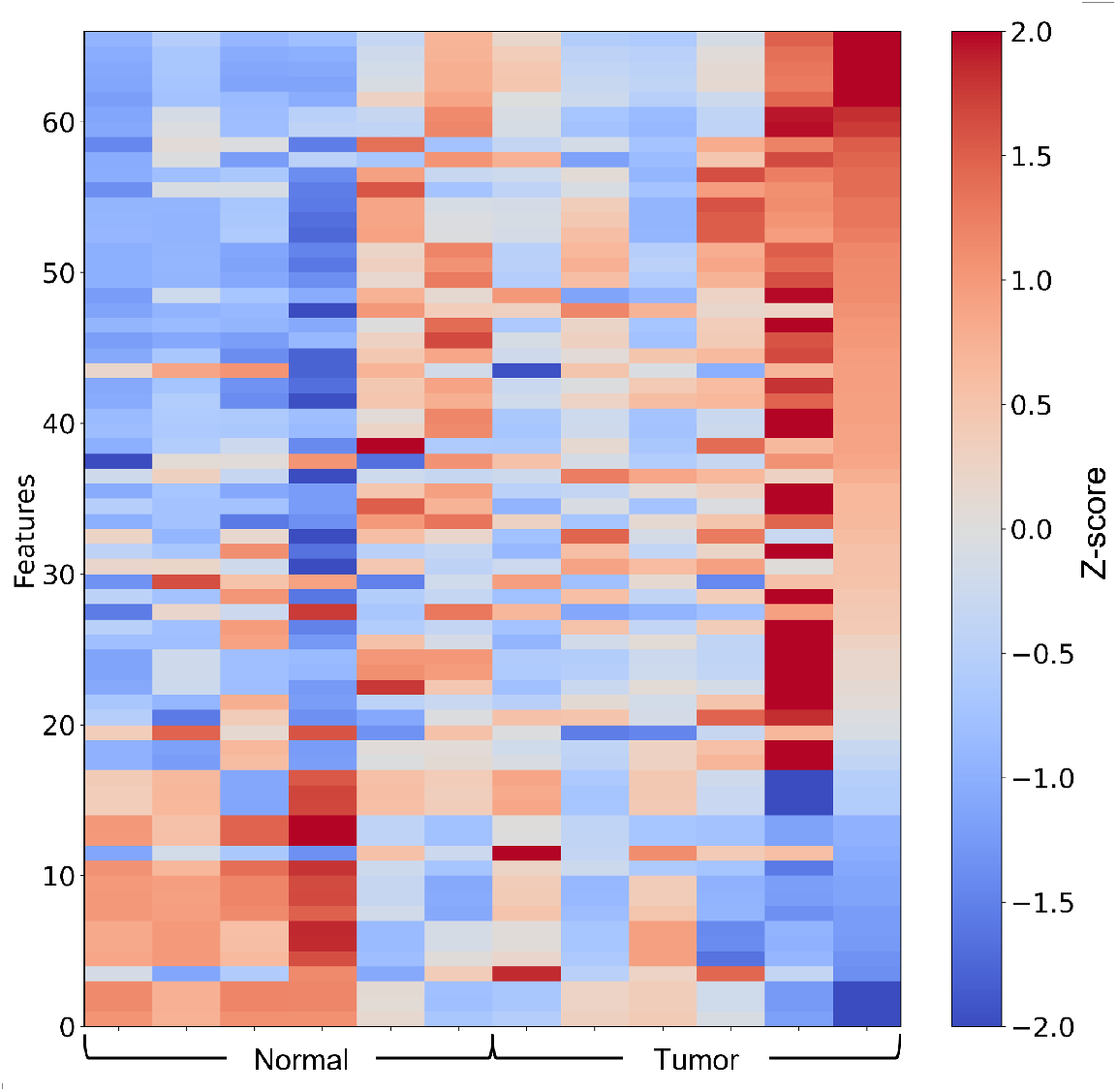
Visualization of features between tumor and normal tissue demonstrated by Z-scores of features for each sample.

### 3.2 Classification

Figure 6 demonstrates LDA results using only the SSTR2 feature. Since there is only one feature, we visualize the two classifiers using a bar chart in Figure 6A. The average intensity is lower in the normal samples compared to the tumor samples as expected, given SSTR2 is overexpressed in most PNETs. However, the ROC curve in Figure 6B shows that LDA is unable to provide high sensitivity between tumor and normal tissue using this singular feature, suggesting that SSTR2 imaging alone could benefit from the inclusion of additional modalities.

**Figure 6.**
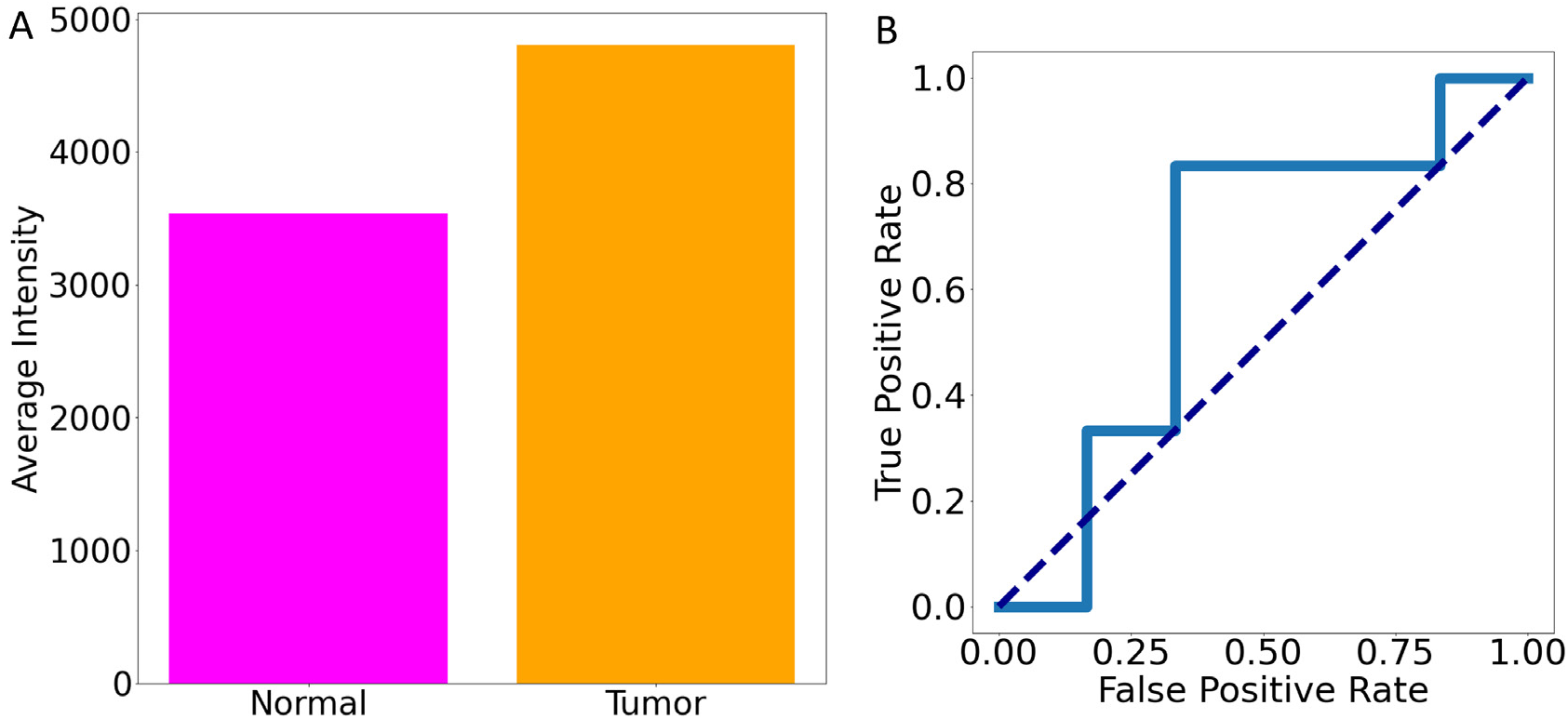
(A) Bar chart of average SSTR2 feature value for tumor and normal tissue. (B) ROC curve for classifiers developed using only the SSTR2 feature.

Table 3 shows LDA accuracy, defined as a simple ratio of correct classifications to total samples, using different numbers of feature sets of both MPM and SSTR2 fluorescence images. Figure 7A shows an LDA projection for n=4 features, and Figure 7B the ROC curves for n=1,2,3,4 features. Accuracy increases as the number of features increases until it reaches 100% at n=4 features. This indicates that MPM can be used to differentiate between diseased and normal tissue to high sensitivity.

**Table 3.**
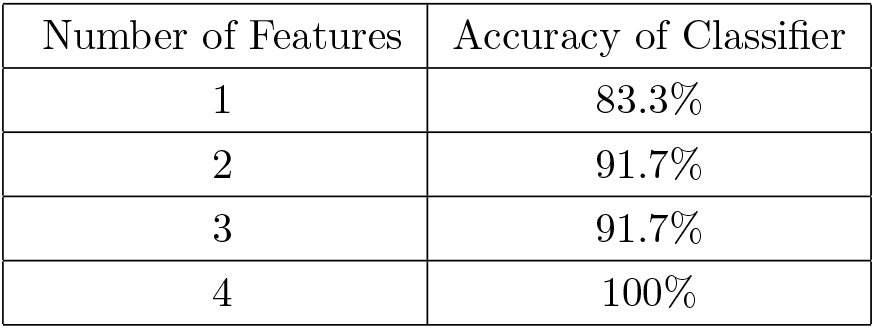
Number of features included in each classifier and average accuracy of the classifier in distinguishing between tumor and normal tissue. Analysis was done using both MPM and SSTR2 fluorescence images.

**Figure 7.**
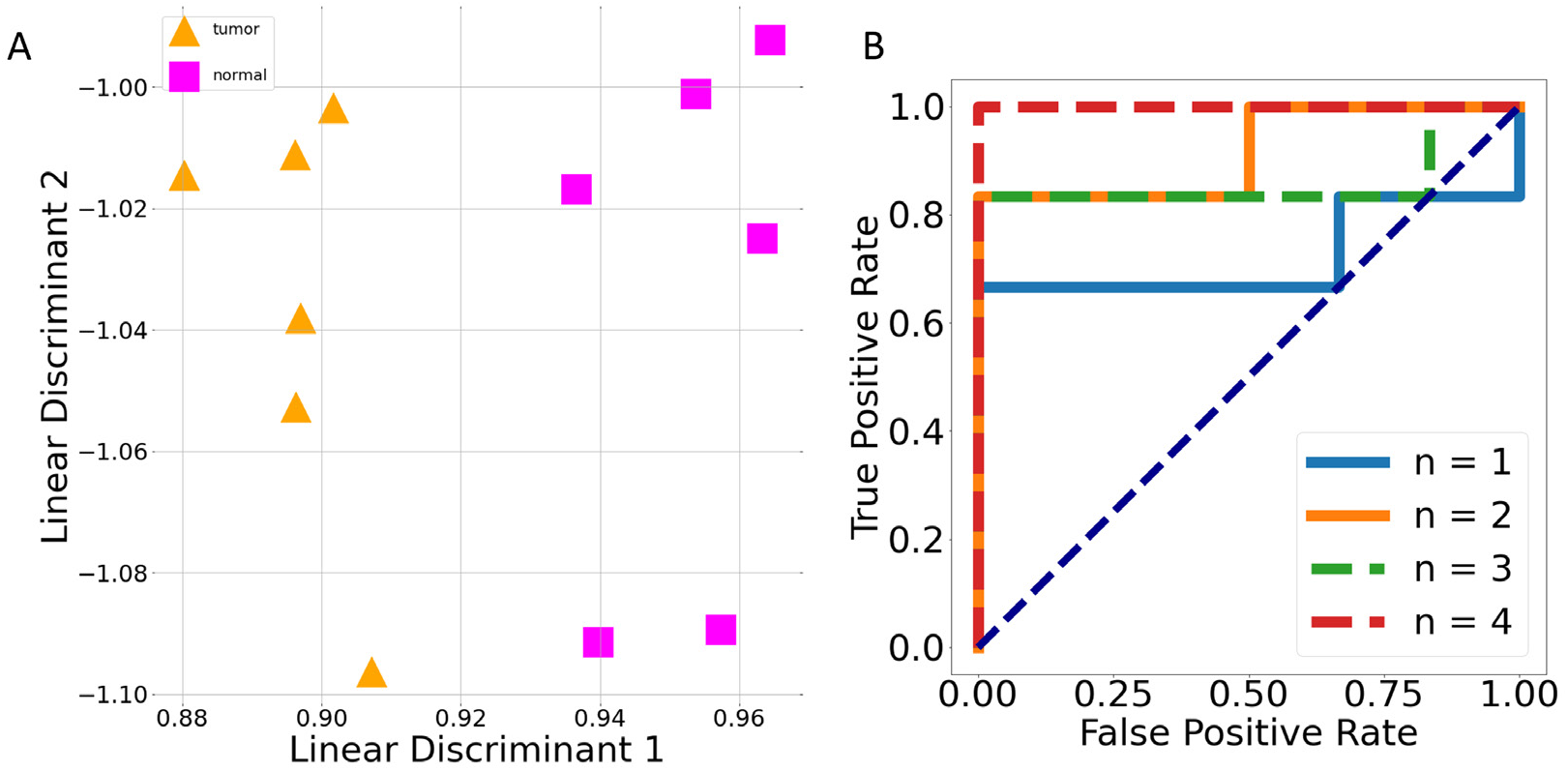
(A) LDA projection using n=4 features to define classifiers and (B) ROC curve for classifiers developed using both MPM and SSTR2 features.

## 4. CONCLUSIONS

In this proceeding, we measure optical characteristics of PNETs and normal pancreatic tissue using MPM and fluorescence imaging of a novel SSTR2-conjugated peptide, and applied linear discriminant analysis to classify the two tissue types. We demonstrate that four texture features are sufficient to classify tumor and normal tissue with 100% accuracy using both MPM and SSTR2 images. Using only MPM images, we can obtain the same accuracy with the same number of features, indicating MPM could be used to determine surgical margins with high sensitivity and specificity. Using only SSTR2 images and one feature, we obtain an accuracy of 66.6%, indicating that SSTR2 provides sensitivity to disease, but the addition of MPM can improve sensitivity and specificity.

## ACKNOWLEDGMENTS

We would like to thank other members of the Merchant and Sawyer laboratories for support, Patricia Jansma at the Imaging Core Marley at the University of Arizona, as well as our collaborators in the Azhdarinia laboratory at the University of Texas Health Science Center for supplying the near infrared SSTR2 dye and collaborators at MD Anderson for clinical expertise and guidance. This work was funded by National Institutes of Health Grant Number R01 DK45729-27, and National Cancer Institute of the National Institutes of Health award number P30 CA023074.

## Notes

### Competing Interest Statement

The authors have declared no competing interest.

